# Referential and attentional accounts of dog point-following in an asymmetric multi-cup design

**DOI:** 10.64898/2026.05.05.722884

**Authors:** Jennifer D. Mugleston, Shin-Miau Huang, Christoph D. Dahl

**Author notes:** Shared authorship (Jennifer D. Mugleston, Shin-Miau Huang).

## Abstract

Human pointing is often used to test whether dogs extract object-specific information from human communicative cues. However, above-chance responses in standard object-choice tasks do not by themselves distinguish between a referential interpretation, in which the gesture identifies a specific target, and an attentional interpretation, in which it primarily biases behaviour toward a broader spatial region. We addressed this issue using an asymmetric six-cup arrangement designed to separate coarse side guidance from exact cup localisation more clearly than a symmetric multi-cup design. Performance in domestic dogs was analysed using three measures: the probability of reaching the correct side, the probability of choosing the correct cup overall, and the probability of choosing the correct cup conditional on having first reached the correct side. The principal comparison involved three matched trial classes: the symmetric 3-vs-3 condition, 2-vs-4 trials with the baited cup on the 2-cup side, and 2-vs-4 trials with the baited cup on the 4-cup side. Descriptively, pointing trials exceeded matched no-point control trials more clearly for side selection than for overall cup choice. The clearest condition effect was observed at the level of side guidance. Dogs were most likely to reach the correct side when the baited cup was located on the 4-cup side of the unequal arrangement. Mixed-effects models confirmed a reliable group effect for side accuracy, whereas overall cup accuracy showed only a weaker and less robust condition effect, and within-side localisation revealed no reliable group difference once condition-specific chance baselines were taken into account. A complementary generative model comparison converged on the same conclusion: a referential-only model fit poorly, an attention-only model captured most of the grouped outcome structure, and a combined model yielded only a modest improvement. Dog point-following is therefore best understood as a layered process dominated by attentional guidance, with only limited additional target-specific localisation.

## Introduction

Domestic dogs (*Canis lupus familiaris*) have lived alongside humans for at least tens of thousands of years, and domestication is widely thought to have favoured traits facilitating interspecific social interaction. Against that background, dogs’ sensitivity to human behavioural and communicative cues has often been treated as a central comparative question (Miklösi et al., 1998; Miklósi and Topál, 2005; Kaminski and Nitzschner, 2013). Human pointing gestures have become a particularly influential test case, because they provide a simple and experimentally tractable way to ask whether dogs can use a human-produced signal to guide behaviour toward a hidden target (Miklösi et al., 1998; Soproni et al., 2002; Virányi et al., 2008; Lakatos et al., 2009). At the same time, the theoretical significance of success in the pointing paradigm remains unresolved. The central issue is not merely whether dogs are affected by pointing, but what kind of information they extract from it. Under a strong referential interpretation, the gesture is taken to indicate a specific object or location, such that the recipient identifies the intended referent (Kaminski and Nitzschner, 2013). Under a weaker attentional interpretation, the gesture primarily biases attention or movement toward a broader spatial region, without robust identification of the exact target (Udell et al., 2010; Dorey et al., 2010; Elgier et al., 2012; Wynne, 2016). This distinction matters because the same above-chance choice pattern can in principle arise from multiple underlying processes, including learned cue use (Udell et al., 2010), directional bias (Dorey et al., 2010), or coarse spatial guidance (Elgier et al., 2012; Wynne, 2016), and therefore does not by itself establish human-like referential understanding. Even authors who take dog point-following seriously as a socio-cognitive phenomenon have emphasized that successful responding does not automatically settle the mechanism question (Kaminski and Nitzschner, 2013). This ambiguity becomes more pressing when one considers the diversity of the existing literature. Performance has been reported to vary with cue type and gesture distance (Dorey et al., 2010), with prior experience (Udell et al., 2013), with ostensive signalling and communicative context (Duranton et al., 2017), and with training history and related procedural factors across studies (Byosiere et al., 2023). Reviews and critical discussions have therefore noted that broad cognitive conclusions are often drawn from paradigms that differ substantially in method, control structure, and inferential leverage (Udell et al., 2010; Kaminski and Nitzschner, 2013; Clark et al., 2019). More recently, the ManyDogs consortium reported only weakly above-chance performance in a standardised two-choice pointing task, reinforcing the need for caution when interpreting small behavioural effects as evidence of robust referential comprehension (ManyDogs Project et al., 2023).

A recent preprint sharpened this issue by modifying the standard pointing task so that dogs were tested not only in a two-choice setting, but also in a six-choice setting (3-vs-3) (Bowers et al., 2025). This increase in the number of alternatives is important because it allows performance at the level of exact cup choice to be separated more clearly from performance at the level of broader directional guidance. Against that background, Bowers et al. (2025) formalized two competing accounts of dog point-following as distinct scaling predictions. Under a referential account, correct choices reflect successful identification of the specific target on some proportion of trials, with the remaining responses attributable to guessing. Under an attentional-capture account, by contrast, the gesture produces a systematic directional bias on otherwise noisy choice behaviour. These two accounts therefore make different quantitative predictions as the number of alternatives increases. In their data, dogs performed only slightly above chance in selecting the correct cup in both the two-choice and six-choice tasks, whereas correct-side performance in the six-choice task was markedly stronger than exact cup performance (Bowers et al., 2025). This pattern was interpreted as more consistent with coarse directional guidance than with strong cup-specific reference.

A closely related question is whether the same contrast can be evaluated directly within the present unequal multi-cup design by fitting simple generative models to the observed outcome counts. Such models make it possible to ask not only whether side- and cup-level performance dissociate descriptively, but also whether the resulting pattern is better captured by a pure attention account, a pure referential account, or a model in which a dominant attentional component is supplemented by a weaker referential contribution. For that reason, the present study combines trial-wise mixed-effects analyses with a complementary group-level model comparison. This logic motivates the present study, but it also highlights a limitation of the standard symmetric six-choice design. A 3-vs-3 arrangement dissociates side selection from exact cup selection, because choosing the correct side is no longer identical to choosing the correct cup. However, both sides remain equally complex. Thus, although a symmetric design can show that side-following and cup-following come apart, it cannot test whether exact cup localisation depends on the number of alternatives remaining within the indicated side after the correct side has been reached. The present analysis addresses this issue through an unequal 2-vs-4 arrangement. In this design, one side of the arena contains two cups and the other contains four. This asymmetry provides stronger diagnostic leverage than a symmetric 3-vs-3 layout. Under a referential account, performance should depend primarily on whether the dog identifies the pointed-to cup, and therefore relatively little on whether that cup lies on a side containing two alternatives or four. Under an attentional account, by contrast, the gesture should first guide the dog toward the correct side or region, after which exact cup choice should depend on the number of alternatives within that side. The unequal arrangement therefore permits a more discriminating decomposition of behaviour into coarse side guidance and within-side localisation than is possible in a symmetric design.

To evaluate these possibilities, performance was decomposed into three quantities: the probability of reaching the correct side, the probability of choosing the correct cup overall, and the probability of choosing the correct cup conditional on having first reached the correct side. This decomposition permits the central question of the study to be addressed directly: whether the unequal side structure primarily alters coarse region-level guidance, precise referent localisation, or both. The present analysis was structured around the following hypotheses.

### Hypothesis 1: coarse side guidance

If dogs use the point primarily as a directional cue, then side accuracy should exceed chance. In a six-cup arena with two sides, chance for side accuracy is *P*(correct side by chance) 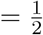.

### Hypothesis 2: weak exact cup localisation

If dogs do not robustly identify the pointed cup as a specific referent, then exact cup accuracy should be much lower than side accuracy, even when overall performance remains above chance.

### Hypothesis 3: side-size dependence under the unequal arrangement

If behaviour follows a side-first attentional process, then within-side localisation should depend on how many alternatives remain once the dog is on the correct side. In the unequal design, chance accuracy conditional on being on the correct side is *P*(correct cup | correct side, 2-side) 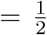 and *P*(correct cup | correct side, 4-side) 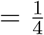. In a symmetric 3-vs-3 design, the corresponding conditional chance level is *P*(correct cup |correct side, 3-side) 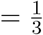.

### Hypothesis 4: dissociation of side and cup effects

If the point primarily shifts behaviour toward a broad spatial region, then the unequal arrangement may generate opposing effects at different levels of analysis. The larger side may be easier to approach as a region, while exact cup localisation within that side may remain difficult.

The aim of the present analysis was therefore not merely to ask whether dogs chose the correct cup above chance, but to decompose performance into side selection and within-side cup selection, and to determine whether the resulting pattern is more consistent with a referential account or an attentional account.

## Methods

### Subjects

The analysed data were obtained from domestic dogs (*Canis lupus familiaris*) tested individually in a point-following paradigm. Dogs participated voluntarily together with their owners. Subject-level information recorded for each dog included training level, training time, walking side, age, sex, breed, FCI group (Fédération Cynologique Internationale), and head shape. To ensure sufficient participation across the full procedure, a completion criterion was imposed. Subjects completing less than 50% of trials were excluded from the analyses reported here. One dog met this exclusion criterion, having completed less than 15% of the procedure. The retained sample therefore comprised 26 dogs, all of which completed the full testing procedure. Subject characteristics for these 26 dogs are summarised in Table 1; the excluded dog is not included in the table.

**Table 1:** Subject characteristics for the analysed dogs. IDs are anonymized. Age is given in years. Training time is reported as an ordered category code: 1 = no training, 2 = less than 3 months of training, 3 = 3 to 12 months of training, and 4 = more than 12 months of training. FCI = Fédération Cynologique Internationale breed group. Breed labels for mixed-breed dogs are descriptive categories; FCI-group entries for these dogs should be interpreted as approximate rather than formal breed classifications.

| ID | Sex | Age<br>(years) | Breed | FCI group | Head shape | Training<br>level | Training time<br>category |
| --- | --- | --- | --- | --- | --- | --- | --- |
| 1 | Female | 3 | Mixed-breed dog | 5 | Meso | 1 | 4 |
| 2 | Female | 2 | Chihuahua | 9 | Brachy | 3 | 3 |
| 3 | Female | 3 | Formosan mixed-breed dog | 5 | Dolicho | 1 | 4 |
| 4 | Male | 8 | Mixed-breed dog | 5 | Meso | 1 | 4 |
| 5 | Male | 5 | Mixed-breed dog | 5 | Meso | 1 | 4 |
| 6 | Male | 11 | Pomeranian-type mix | 5 | Meso | 0 | 1 |
| 7 | Female | 15 | Formosan mixed-breed dog | 5 | Meso | 0 | 1 |
| 8 | Male | 6 | Mixed-breed dog | 1 | Meso | 1 | 2 |
| 9 | Female | 7 | Wire Fox Terrier | 3 | Meso | 3 | 4 |
| 10 | Female | 4 | Hound-type mix | 6 | Meso | 2 | 4 |
| 11 | Male | 9 | Poodle-type mix | 9 | Meso | 2 | 3 |
| 12 | Male | 3 | Shiba Inu | 5 | Meso | 1 | 4 |
| 13 | Male | 6 | Border Collie | 1 | Dolicho | 1 | 3 |
| 14 | Male | 2 | Shiba Inu | 5 | Meso | 2 | 3 |
| 15 | Female | 1 | Yorkshire Terrier | 3 | Meso | 2 | 3 |
| 16 | Male | 1 | Australian Shepherd | 1 | Meso | 2 | 3 |
| 17 | Male | 2 | Labrador Retriever | 8 | Meso | 1 | 1 |
| 18 | Female | 10 | Shiba-type mix | 5 | Meso | 1 | 2 |
| 19 | Female | 3 | Border Collie | 1 | Meso | 2 | 4 |
| 20 | Female | 6 | Japanese Spitz | 5 | Meso | 1 | 2 |
| 21 | Female | 3 | Schnauzer | 2 | Meso | 2 | 4 |
| 22 | Male | 5 | Herding-type mix | 1 | Meso | 1 | 4 |
| 23 | Female | 3 | Standard Poodle | 9 | Dolicho | 2 | 4 |
| 24 | Male | 5 | Spaniel-type dog | 9 | Meso | 1 | 2 |
| 25 | Female | 4 | Formosan mixed-breed dog | 5 | Dolicho | 1 | 2 |
| 26 | Female | 5 | Dobermann | 2 | Dolicho | 1 | 2 |

**Table 2:** Interpretive logic of the present decomposition. “Side” refers to whether the dog reaches the correct side of the arena. “Cup” refers to whether, once on the correct side, the dog selects the correct cup above the relevant within-side chance level.

| Side | Cup | Key idea | Interpretation |
| --- | --- | --- | --- |
| Low | Low | No usable cue effect | No clear evidence that the pointing gesture guides behaviour. Responses are consistent with random choice or failure to extract useful information from the cue. |
| High | Low | Side only | Most consistent with **attention / directional guidance**. The dog reaches the correct side, but does not identify the correct cup more precisely than expected by within-side chance. |
| Low | High | Unstable pattern | Difficult to interpret. This would imply poor side guidance but unexpectedly precise cup selection once on the correct side. In practice, such a pattern would require careful scrutiny. |
| High | High | Side + cup | Most consistent with **referential localisation**. The gesture appears to guide the dog both toward the correct side and toward the specific target cup. |
| High on large side | Weak / inconsistent cup effect | Region attraction | Suggests a **broad spatial bias**. The larger side may function as a stronger attractor, improving region entry without producing equally strong exact cup localisation. |

### Ethical Note

The study was non-invasive and involved privately owned domestic dogs participating voluntarily with their owners. Owners provided informed consent before participation and remained present throughout testing. Dogs were not restrained beyond being held briefly by the owner at the marked starting position before release, and they were free not to respond on any trial. The task involved food rewards and standard object-choice procedures, with no aversive stimulation, deprivation, or physical manipulation of the dogs. If a dog did not make a valid choice within the response window or otherwise failed to complete a trial, the trial was recorded as a no-response or incomplete trial, and the dog was called back by the owner for the next trial. One dog that completed less than 15% of the procedure was excluded from analysis under the predefined completion criterion. The procedure was designed to minimise stress and to keep owner–dog interaction under normal handling conditions.

### Task

The task was conducted in a six-cup choice setup. The spatial layout of the six-cup arrangement, including the relative positions of the dog, experimenter, and cups, is illustrated in Figure 1A. The cups measured approximately 10 × 10 × 7.5 cm and were arranged on a semicircular arc around the dog. Each cup was positioned approximately 1.5 m from the starting location, with approximately 32 cm spacing between neighbouring cups. The dog sat at the centre of the arc on a marked starting location indicated by a pet pad and faced the experimenter along the central axis of the arrangement. The owner held the dog at this location during the manipulation and cue-presentation phases of the trial. The experimenter stood on the central axis between the two innermost cups during cue presentation and then took a small step backwards after pointing, so that the dog was encouraged to approach the cup array rather than the experimenter. In the symmetric condition, the cups were arranged in a 3-vs-3 layout, with three cups on each side of the central axis. In the unequal conditions, the six cups were distributed asymmetrically as either a 2-vs-4 or a 4-vs-2 layout. These labels indicate the number of cups on the left and right sides, respectively. Thus, in a 2-vs-4 trial, two cups were located on the left side and four on the right side, whereas the reverse was true in a 4-vs-2 trial. These two labels refer to mirror-image spatial layouts and were pooled at the level of the main manipulation because both instantiate the same structural asymmetry, namely that one side contains two alternatives and the other contains four. However, cue position was retained trial by trial so that it could be determined whether the pointed-to cup lay on the 2-cup side or on the 4-cup side.

**Figure 1:**
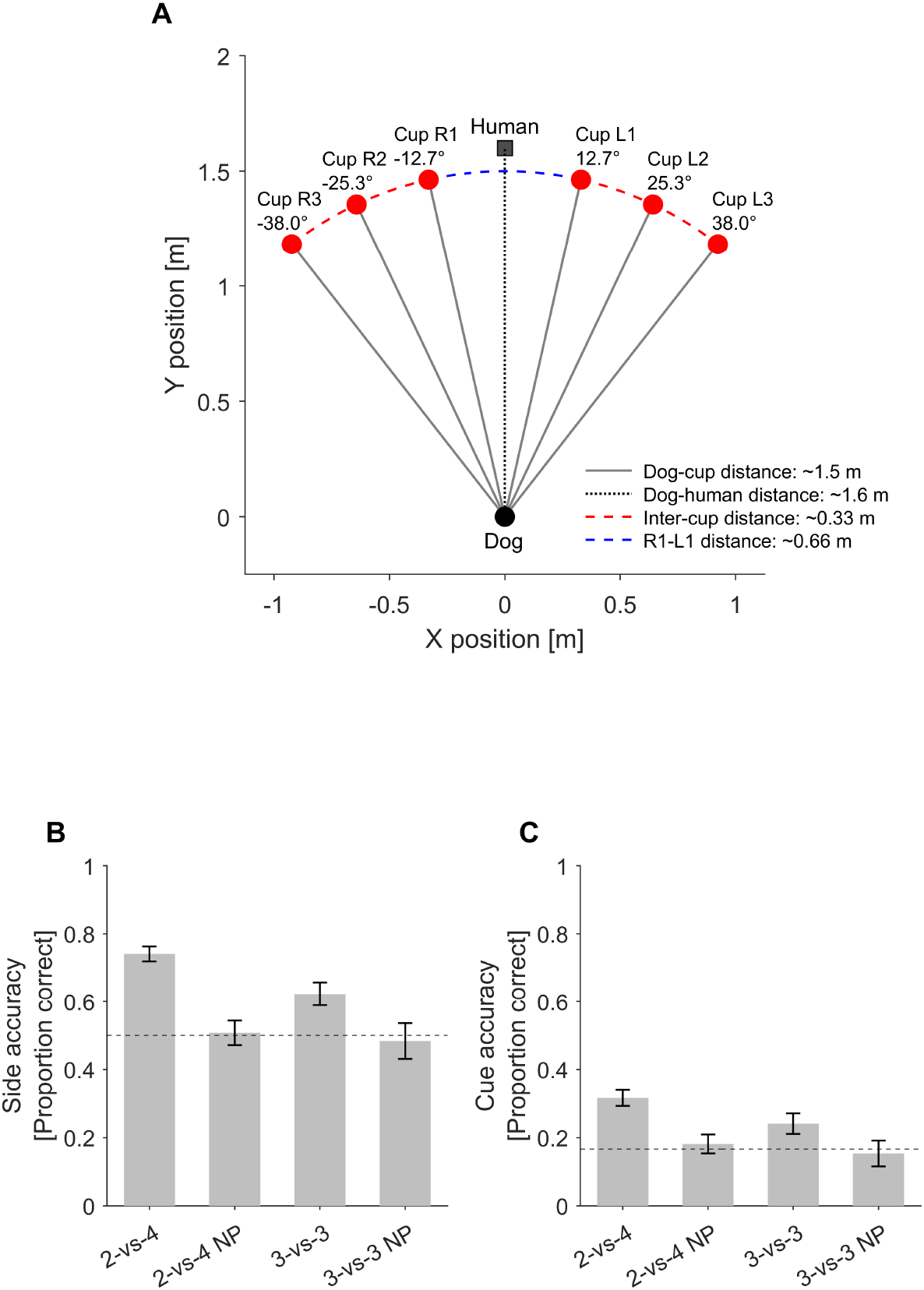
Task geometry and descriptive comparison of pointing and no-pointing conditions. Panel A illustrates the six-cup arrangement used in the experiment, including the dog position, the central human position, and the spatial layout of the cups relative to the dog. Panels B and C show descriptive performance across the pooled 2-vs-4 and 3-vs-3 arrangements, with corresponding no-point controls. Panel B shows side accuracy; the dashed horizontal line indicates side-level chance performance (0.5). Panel C shows overall cue accuracy; the dashed horizontal line indicates overall chance performance in the six-cup task (1/6). Pointing improved side-level performance more clearly than overall cup-level performance, and the clearest descriptive separation from the no-point controls was observed in side accuracy for the pooled 2-vs-4 arrangement.

At the start of each trial, the experimenter manipulated all cups sequentially while the dog observed the procedure. To standardise handling and prevent the dog from identifying the baited cup directly, each cup was covered in turn with a 30 × 30 cm panel while the experimenter acted behind it. From the dog’s perspective, all cups were therefore manipulated in the same general way, and the baited cup could not be identified simply from visible hand movements. To reduce the possibility that dogs could solve the task using olfactory cues, each cup contained a small bait stick attached to the upper inside. The experimentally baited cup therefore did not stand out as the only cup carrying food odour. Instead, any local food smell was diluted against a common background odour across the array. This same manipulation procedure was used in both pointing and no-point control trials. The critical difference was that, in the no-point trials, the experimenter did not provide the subsequent pointing cue. The communicative cue then followed a fixed sequence. The experimenter first called the dog’s name to ensure attention. Once the dog was attending, the experimenter briefly shifted gaze toward the target cup and then pointed to that cup for approximately 2 s. After pointing, the arm was returned behind the back, where the other arm was already held. During the pointing interval, the experimenter maintained eye contact with the dog except for the initial brief gaze toward the cup. Immediately after the pointing gesture, the experimenter maintained a straight-ahead gaze directed beyond the dog rather than at its face, in order to minimize the possibility that the dog would approach the experimenter instead of the indicated cup location. Following the verbal go signal given by the experimenter, the owner released the dog from the marked starting position, allowing it to approach one of the cups. A choice was defined by the dog entering a marked area in front of one of the cups. These marked areas served as predefined choice zones and measured approximately 30 × 20 cm. If the dog did not enter any choice zone within 20 s after release, the trial was recorded as a no-response trial. If the dog chose the baited cup, it received the reward. If the dog chose incorrectly, the owner called the dog back in preparation for the next trial. In addition to pointing trials, no-point control trials were available for the broader condition summaries (3-vs-3 no-point, 2-vs-4 no-point, and 4-vs-2 no-point). These trials preserved the same spatial arrangements and manipulation procedure but omitted the pointing gesture, thereby providing a baseline for spontaneous choice behaviour in the absence of the communicative cue. Warm-up and training trials were marked separately in the trial records and were excluded from the present analyses. These comprised the successive warm-up phases 1-vs-1, 3-vs-3, 4-vs-2, and 2-vs-4, followed by the no-point variants of the unequal arrangements.

For each analysed trial, two binary outcome variables were derived. The first, `side_correct`, indicated whether the dog first moved to the correct side of the arena. The second, `cue_correct`, indicated whether the dog chose the baited cup. This distinction allowed behaviour to be decomposed into side-level guidance and exact cup localisation. In the unequal arrangement, this decomposition was especially informative because the within-side chance level differs depending on which side was cued: *P*(correct cup | correct side, 2-side) 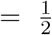and *P*(correct cup| correct side, 4-side) 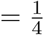. In the symmetric 3-vs-3 arrangement, the corresponding conditional chance level is *P*(correct cup |correct side, 3-side) 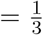. These condition-specific baselines made it possible to assess not only whether dogs reached the correct side, but also how precisely they localised the target cup once the correct side had been reached.

### Data preprocessing and analysis

The data were organised at the trial level according to subject identity, condition, cue position, block, and the two derived binary outcome variables `side_correct` and `cue_correct`. Trials recorded as no-response trials were retained in the dataset but coded as missing for `side_correct` and `cue_correct`, because no valid side choice or cup choice had been made. Accordingly, they contributed to trial counts and condition summaries of participation, but not to the binomial outcome models for side accuracy or cup accuracy. The principal analytical comparison was structured around three trial classes: the symmetric 3-vs-3 condition, unequal-arrangement trials (2-vs-4 / 4-vs-2) in which the baited cup was located on the 2-cup side, and unequal-arrangement trials in which the baited cup was located on the 4-cup side. For trials from the unequal arrangement, cue position was combined with the raw condition label to determine whether the cued cup lay on the 2-cup side or on the 4-cup side. These classes differed somewhat in trial count after exclusions and no-response variation, but the mixed-effects models used here accommodate such imbalance directly. This distinction was critical because the relevant within-side chance baseline differs across these cases. The resulting trial structure allowed performance to be analysed hierarchically. Side-level performance was quantified as the probability that the dog first reached the correct side of the arena. Overall cup-level performance was quantified as the probability that the dog selected the baited cup. Because selecting the correct cup necessarily implies being on the correct side, exact within-side localisation was quantified as the probability of selecting the baited cup conditional on having first reached the correct side. In this formulation, overall cup success can be decomposed into the probability of reaching the correct side and the probability of identifying the correct cup once that side has been reached. Condition-specific within-side chance baselines were therefore 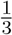 for the symmetric 3-vs-3 condition,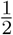 for unequal-arrangement trials in which the baited cup was located on the 2-cup side, and 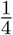 for unequal-arrangement trials in which the baited cup was located on the 4-cup side. This made it possible to distinguish coarse side guidance from exact cup localisation. For each of the three trial classes, descriptive summaries were computed for side accuracy and overall cup accuracy as means with standard errors. In addition, conditional cup accuracy was calculated using only those trials on which the dog had first reached the correct side. This yielded a condition-wise estimate of the precision with which dogs selected the correct cup after already having entered the correct side of the arena.

To evaluate the effects of the three trial classes on side accuracy, overall cup accuracy, and within-side localisation while accounting for repeated observations within subjects, trial-wise binomial models with dog identity as a subject-level random factor were fitted separately for the three outcome measures. The first model examined side accuracy as the dependent variable. The second examined overall cup accuracy. The third was restricted to trials on which the dog had already reached the correct side and examined exact cup localisation conditional on correct-side entry. In this third analysis, the relevant condition-specific chance baselines were incorporated directly so that the model tested deviations from the expected within-side chance level for each trial class. In addition to the main condition factor, supplementary trial-level predictors were considered to assess whether the observed effects might depend on simpler spatial or procedural features. These predictors comprised cue side (left vs. right), cue position number, and block. This permitted evaluation of whether the principal effects of interest were attributable to the structure of the arrangement itself or instead reflected lateral bias, positional effects, or changes across successive portions of the session. As an exploratory supplementary analysis, additional mixed-effects models were also fitted including selected dog-level metadata variables. Candidate metadata variables comprised age, training level, training time, sex, head shape, FCI group, and walking side. Training time was treated as an ordered coded predictor, with higher values corresponding to longer training history. Because of sparsity and model-stability considerations, reduced supplementary models were fitted using selected subsets of these predictors for the different outcome measures. Their inclusion was intended to assess whether individual differences in subject characteristics contributed to side accuracy, overall cup accuracy, or within-side localisation beyond the principal task predictors.

### Generative modelling of outcome counts

In addition to the trial-wise mixed-effects analyses, a grouped model-comparison approach was used to evaluate three simple process accounts of performance: an attention-only model, a referential-only model, and a combined model. These models were fitted to the grouped counts for the three analytically critical pointing subsets: the symmetric `3vs3` condition, the `2vs4` subset with the baited cup on the 2-cup side (`2vs4_2side`), and the `2vs4` subset with the baited cup on the 4-cup side (`2vs4_4side`). For each model, predicted probabilities were derived for side accuracy, overall cup accuracy, and conditional cup accuracy given correct-side entry. Model fit was evaluated by log-likelihood, Akaike’s information criterion (AIC), and the Bayesian information criterion (BIC), and the resulting predictions were compared directly with the observed values. For each of the three analytically critical pointing subsets, trials were classified into three mutually exclusive outcome categories: correct cup choice, incorrect cup choice on the correct side, and incorrect choice on the wrong side. These grouped counts preserve the central behavioural distinction between coarse side guidance and exact cup localisation while allowing simple mechanistic accounts to be formalized explicitly. Three models were compared. In the *attention-only* model, a single parameter *p*_*g*_ for each group *g* specifies the probability that the dog is guided to the correct side. Once on that side, choice among the cups on that side is random. If *k*_*g*_ denotes the number of cups on the correct side (*k*_*g*_ ∈{3, 2, 4} for `3vs3, 2vs4_2side`, and `2vs4_4side`, respectively), then

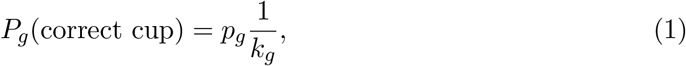

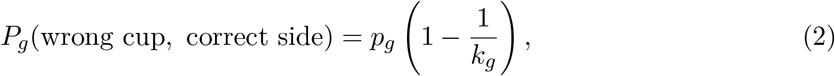

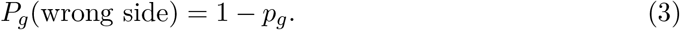

Thus, the model predicts *P*_*g*_(correct side) = *p*_*g*_ and fixes conditional cup accuracy at the relevant within-side chance level. In the *referential-only* model, a single parameter *q*_*g*_ specifies the probability that the dog identifies the exact target cup directly. On the remaining trials, choice is assumed to be random across all six cups. The resulting outcome probabilities are

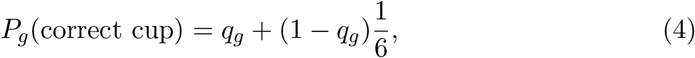

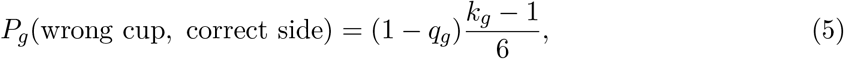

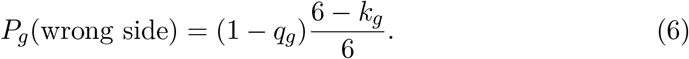

This model therefore predicts both side and cup performance from exact target identification plus global guessing. In the *combined* model, exact referential identification occurs with probability *q*_*g*_, whereas on the remaining trials behaviour follows the attention process with probability *p*_*g*_. The resulting outcome probabilities are

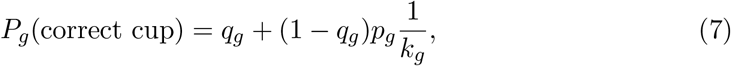

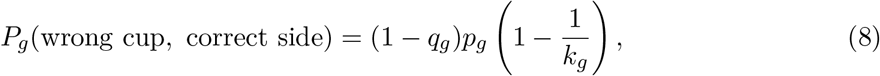

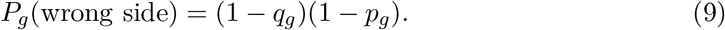

Accordingly, this model predicts

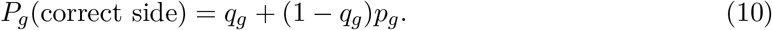

Parameters were estimated by maximum likelihood independently for each group from the multinomial outcome counts, and the resulting log-likelihoods were summed across groups for model comparison. Because the pure attention and pure referential models each contain one free parameter per group, both have three free parameters in total, whereas the combined model contains six.

## Results

### Condition-wise summaries

The descriptive results are summarised in Figures 1 and 2. Figure 1B–C compare pointing and no-pointing performance across the pooled 2-vs-4 and 3-vs-3 conditions, separately for side accuracy and cue accuracy. Figure 2 then decomposes performance within the asymmetric arrangement into side accuracy, overall cup accuracy, and conditional cup accuracy as a function of whether the baited cup was located on the 2-cup side or the 4-cup side. At the broader condition level, pointing conditions showed higher descriptive performance than no-point controls, especially for side accuracy. In the pooled 2-vs-4 arrangement, side accuracy was 0.741 (*SE* = 0.022) with pointing and 0.508 (*SE* = 0.036) in the no-point control. In the symmetric 3-vs-3 arrangement, the corresponding values were 0.623 (*SE* = 0.034) and 0.484 (*SE* = 0.053). Overall cup accuracy showed a smaller separation: 0.317 (*SE* = 0.023) versus 0.181 (*SE* = 0.028) for the pooled 2-vs-4 arrangement, and 0.241 (*SE* = 0.030) versus 0.154 (*SE* = 0.038) for 3-vs-3. Within the pointing conditions, the asymmetric decomposition yielded the following values:

**Figure 2:**
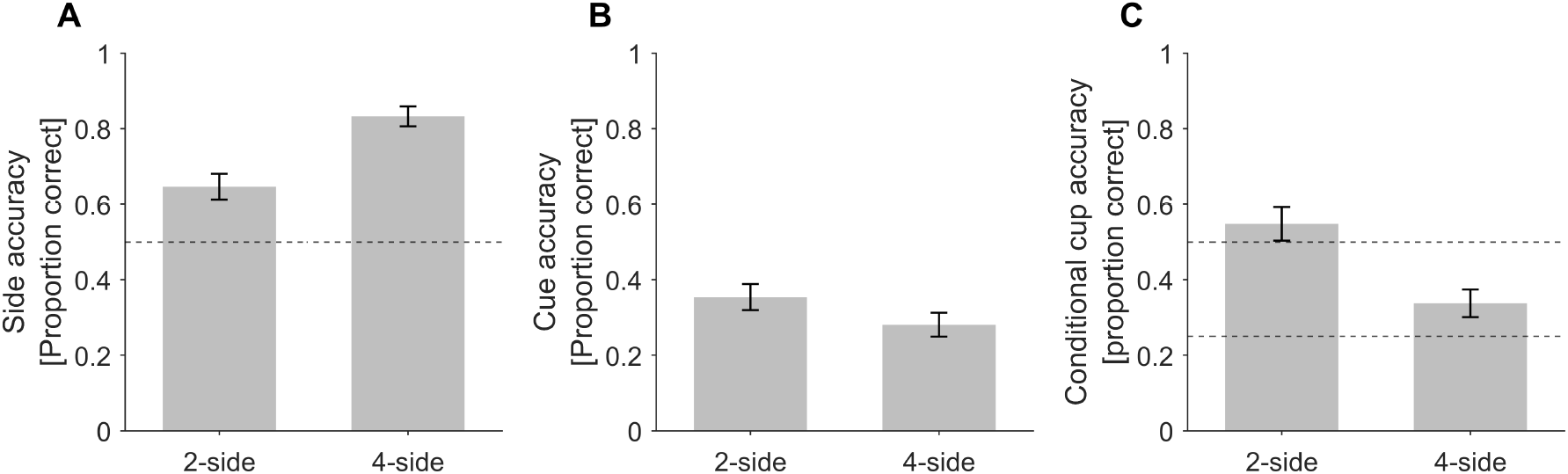
Decomposition of performance within the asymmetric arrangement. Panel A shows side accuracy as a function of whether the baited cup was located on the 2-cup side or the 4-cup side. The dashed horizontal line indicates side-level chance performance (0.5). Panel B shows overall cue accuracy for the same two subsets. Panel C shows conditional cup accuracy given correct-side entry. Dashed horizontal lines in Panel C indicate the relevant within-side chance baselines for the 2-side (1/2) and the 4-side (1/4). The figure separates coarse side guidance from exact cup localisation and shows that the strongest effect of the asymmetric arrangement was expressed at the level of side entry rather than in robust condition differences in conditional cup selection.

### Side accuracy

As shown in Figure 1B and Figure 2A, side accuracy was above chance in all pointing conditions and was highest when the cued cup was on the 4-cup side of the unequal arrangement, with *P*(side_correct |2vs4_4side) = 0.833. By contrast, side accuracy was lower both when the cued cup was on the 2-cup side, with *P*(side_correct |2vs4_2side) = 0.646, and in the symmetric 3-vs-3 condition, with *P*(side_correct| 3vs3) = 0.623. Thus, dogs were especially likely to move to the correct side when the correct side was also the larger side. These values already reveal that side guidance and exact cup localisation did not behave identically. A model-based comparison of attention-only, referential-only, and combined accounts is shown in Figure 3 and is reported below.

**Figure 3:**
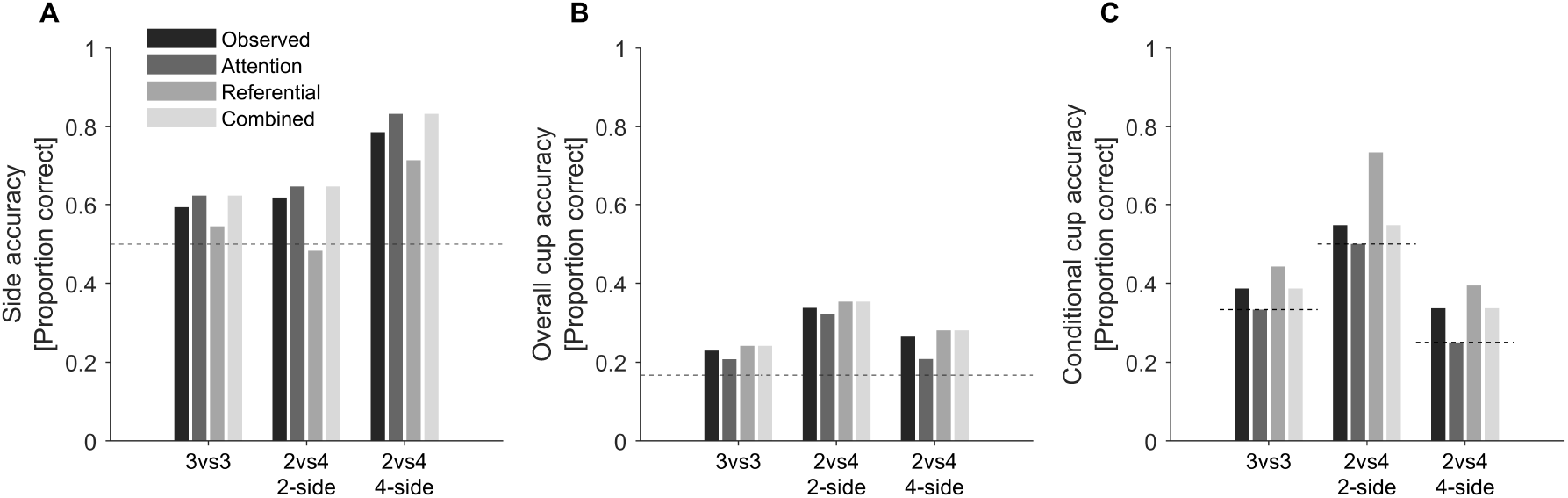
Comparison of observed performance with fitted model predictions. Panel A shows side accuracy, Panel B overall cup accuracy, and Panel C conditional cup accuracy given correct-side entry. In each panel, observed values are shown alongside predictions from the attention-only, referential-only, and combined models for the three analysed trial classes (3vs3, 2vs4 with the baited cup on the 2-cup side, and 2vs4 with the baited cup on the 4-cup side). Dashed horizontal lines indicate the relevant chance baselines: 0.5 for side accuracy, 1/6 for overall cup accuracy, and 1/3, 1/2, and 1/4 for the conditional analyses of the 3vs3, 2-side, and 4-side subsets, respectively. The figure shows that the attention-only model captures the main side-guidance pattern particularly well, whereas the referential-only model overpredicts conditional localisation on the 2-side. The combined model provides the closest overall descriptive match, but with only modest improvement over the attention-only account.

### Overall cup accuracy

As shown in Figure 1C and Figure 2B, overall cup accuracy remained substantially lower than side accuracy. It was somewhat higher for unequal trials in which the cued cup was on the 2-side, with *P*(cue_correct |2vs4_2side) = 0.354. Lower values were obtained both for unequal trials in which the cued cup was on the 4-side, with *P*(cue_correct| 2vs4_4side) = 0.281, and for the symmetric condition, with *P*(cue_correct| 3vs3) = 0.241. Taken alone, these overall cup rates mix two distinct sources of success and failure: whether the dog reached the correct side, and whether it identified the correct cup within that side.

### Conditional cup accuracy given correct side

Once analysis was restricted to trials on which the dog first reached the correct side, a more informative pattern emerged (Figure 2C). When the cue indicated the 2-cup side in the unequal arrangement, conditional cup accuracy was *P*(cue_correct| side_correct, 2-side) = 0.548. This was descriptively above the corresponding within-side chance rate of 0.5, but only by a small margin. When the cue indicated the 4-cup side, conditional cup accuracy was *P*(cue_correct |side_correct, 4-side) = 0.337. This exceeded the corresponding within-side chance rate of 0.25. In the symmetric condition, conditional cup accuracy was *P*(cue_correct |side_correct, 3vs3) = 0.387, which was above the within-side chance rate of 1/3. Thus, exact cup localisation after correct side entry remained limited overall, even though descriptive conditional performance exceeded the relevant within-side chance baselines in all three pointing subsets. The unequal arrangement nevertheless produced an asymmetry in which side selection was strongest when the correct side was large, whereas within-side localisation showed no comparably strong advantage for the 2-side.

### Mixed-effects analysis of side guidance and within-side localisation

To test whether the unequal side structure affected coarse directional guidance and exact cup localisation differently, binomial generalized linear mixed-effects models were fitted at the trial level with dog identity included as a random intercept. The fixed-effect factor `group` distinguished the symmetric condition (`3vs3`) from the unequal-condition subsets in which the cued cup lay on the 2-cup side (`2vs4_2side`) or the 4-cup side (`2vs4_4side`). For the within-side localisation analysis, only trials with `side_correct = 1` were retained, and a group-specific offset was included to account for the relevant conditional chance baselines (1/3, 1/2, and 1/4, respectively). The descriptive patterns motivating these models are visualised in Figures 1 and 2. The full trial-level dataset comprised *N* = 597 observations. The within-side localisation analysis was based on the subset of *N* = 419 trials on which the dog first reached the correct side, comprising 124 trials in `3vs3`, 126 trials in `2vs4_2side`, and 169 trials in `2vs4_4side`.

#### Side accuracy

A mixed-effects model with `side_correct` as outcome showed a significant overall effect of group, *F*(2, 594) = 12.04, *p <* .001 (Table 3). Relative to the `3vs3` reference level, the `2vs4_4side` condition showed a strong positive coefficient (*β* = 1.102, *SE* = 0.239, *p <* .001), whereas the `2vs4_2side` condition did not differ from the reference (*β* = 0.100, *SE* = 0.210, *P*= .635). Thus, dogs were more likely to move to the correct side when the correct side was the 4-cup side of the unequal arrangement. When `cue_lr`, `cue_num`, and `block` were added as additional fixed effects, the group effect remained significant, *F*(2, 586) = 7.69, *P*= .001, and the `2vs4_4side` coefficient remained positive and significant (*β* = 1.133, *SE* = 0.289, *p <* .001). Neither `cue_lr` nor `cue_num` was significant, and the omnibus effect of `block` was not significant.

**Table 3:**
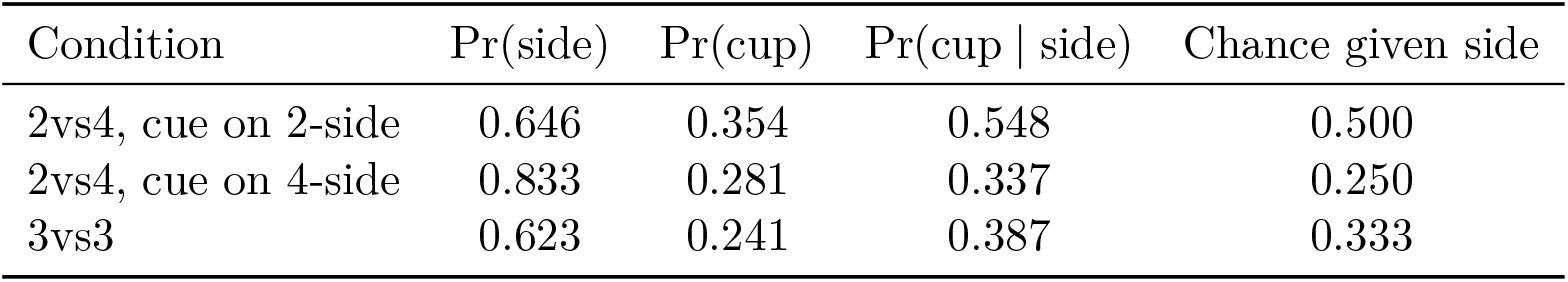
Descriptive performance in the three analytically critical pointing subsets. Reported values are side accuracy, overall cup accuracy, and cup accuracy conditional on correct-side entry, together with the relevant within-side chance baseline for each condition.

**Table 4:** Summary of generalized linear mixed-effects models. Dog identity was included as a random intercept in all models. For the within-side model, the logit-transformed condition-specific chance baseline was included as an offset. The reference level for group was 3vs3.

| Outcome | Effect | Estimate ($\beta$) | SE | Test statistic | $p$ |
| --- | --- | --- | --- | --- | --- |
| Side accuracy | group (omnibus) | — | — | $F(2, 594) = 12.04$ | $< .001$ |
| Side accuracy | 2vs4_2side | 0.100 | 0.210 | $t = 0.474$ | .635 |
| Side accuracy | 2vs4_4side | 1.102 | 0.239 | $t = 4.616$ | $< .001$ |
| Side accuracy + covariates | group (omnibus) | — | — | $F(2, 586) = 7.69$ | .001 |
| Side accuracy + covariates | 2vs4_2side | 0.095 | 0.231 | $t = 0.412$ | .680 |
| Side accuracy + covariates | 2vs4_4side | 1.133 | 0.289 | $t = 3.923$ | $< .001$ |
| Overall cup accuracy | group (omnibus) | — | — | $F(2, 594) = 3.08$ | .047 |
| Overall cup accuracy | 2vs4_2side | 0.544 | 0.223 | $t = 2.435$ | .015 |
| Overall cup accuracy | 2vs4_4side | 0.205 | 0.228 | $t = 0.902$ | .367 |
| Overall cup accuracy + co-<br>variates | group (omnibus) | — | — | $F(2, 586) = 2.18$ | .114 |
| Overall cup accuracy + co-<br>variates | 2vs4_2side | 0.414 | 0.241 | $t = 1.716$ | .087 |
| Overall cup accuracy + co-<br>variates | 2vs4_4side | 0.426 | 0.276 | $t = 1.544$ | .123 |
| Within-side localisation | group (omnibus) | — | — | $F(2, 416) = 0.53$ | .587 |
| Within-side localisation | 2vs4_2side | -0.043 | 0.257 | $t = -0.166$ | .869 |
| Within-side localisation | 2vs4_4side | 0.190 | 0.246 | $t = 0.771$ | .441 |
| Within-side localisation +<br>covariates | group (omnibus) | — | — | $F(2, 408) = 1.90$ | .150 |
| Within-side localisation +<br>covariates | 2vs4_2side | -0.219 | 0.276 | $t = -0.794$ | .428 |
| Within-side localisation +<br>covariates | 2vs4_4side | 0.464 | 0.304 | $t = 1.526$ | .128 |
| Within-side localisation +<br>random slope | group (omnibus) | — | — | $F(2, 408) = 1.59$ | .205 |
| Within-side localisation +<br>random slope | 2vs4_2side | -0.217 | 0.284 | $t = -0.765$ | .445 |
| Within-side localisation +<br>random slope | 2vs4_4side | 0.487 | 0.360 | $t = 1.355$ | .176 |

#### Overall cup accuracy

A corresponding model for `cue_correct` revealed a significant overall effect of group, *F*(2, 594) = 3.08, *p* = .047 (Table 3). Relative to `3vs3`, the `2vs4_2side` condition showed a positive coefficient (*β* = 0.544, *SE* = 0.223, *p* = .015), whereas the `2vs4_4side` condition did not differ significantly from the reference (*β* = 0.205, *SE* = 0.228, *p* = .367). Thus, in the simpler model, overall cup accuracy was elevated specifically for trials in which the cued cup was on the 2-cup side. This pattern weakened after the addition of `cue_lr`, `cue_num`, and `block`. In the extended model, the omnibus effect of group was no longer significant, *F*(2, 586) = 2.18, *p* = .114. Neither `cue_lr` nor `cue_num` was significant, and the omnibus effect of `block` was not significant. Accordingly, the group effect for overall cup accuracy was less robust than the corresponding effect for side accuracy.

#### Within-side localisation

The critical analysis concerned exact cup localisation conditional on the dog having already reached the correct side. In this model, `cue_correct` was analysed on the subset of correct-side trials, and the logit of the condition-specific chance baseline was included as an offset. In the core model, no reliable effect of group was observed, *F*(2, 416) = 0.53, *p* = .587 (Table 3). Relative to `3vs3`, neither `2vs4_2side` (*β* =− 0.043, *SE* = 0.257, *p* = .869) nor `2vs4_4side` (*β* = 0.190, *SE* = 0.246, *p* = .441) differed significantly. The same pattern remained in the extended model including `cue_lr`, `cue_num`, and `block`. The omnibus effect of group remained non-significant, *F*(2, 408) = 1.90, *p* = .150, and in the random-slope version it again remained non-significant, *F*(2, 408) = 1.59, *p* = .205. The coefficient for `cue_num` was negative in all extended models (*p* = .080 and *p* = .064, respectively), suggesting a weak tendency for exact localisation to decline with position number, but this effect did not reach conventional significance.

Taken together, the mixed-effects models indicate that the clearest and most robust condition effect was expressed at the level of side guidance. Overall cup accuracy showed some evidence of condition dependence in the simpler model, driven by the 2-side subset, but this effect weakened after inclusion of cue side, cue position number, and block. By contrast, the unequal arrangement did not produce a reliable change in within-side identification of the target cup.

Exploratory supplementary models including selected dog-level metadata variables did not alter the principal task-related interpretation, but they revealed a selective association between training time and cup-level performance. In the reduced combined metadata models, coded training time was not associated with side accuracy, *β* = 0.045, *SE* = 0.127, *p* = .725. By contrast, longer training history was positively associated with overall cup accuracy, *β* = 0.210, *SE* = 0.104, *p* = .044, and with within-side localisation, *β* = 0.233, *SE* = 0.115, *p* = .044. Thus, training history appeared to modulate the more precise cup-level components of performance rather than the coarse side-guidance component. This secondary individual-difference effect does not change the main conclusion that the clearest task-related condition effect was expressed at the level of side guidance.

### Model-based comparison of attentional and referential accounts

To complement the descriptive decomposition and the mixed-effects analyses, three simple process models were fitted to the grouped behavioural data: an attention-only model, a referential-only model, and a combined model containing both components. Figure 3 summarizes the observed probabilities and the corresponding fitted predictions for side accuracy, overall cup accuracy, and conditional cup accuracy. At the level of side accuracy (Figure 3A), the attention-only model closely reproduced the main empirical pattern, including the particularly high performance in the 2vs4 condition when the baited cup was located on the 4-cup side. The referential-only model captured this pattern less well, especially because it did not reproduce the strong asymmetry in side guidance as effectively as the attention-based account. The combined model also tracked the observed side pattern closely, but did so with only a modest improvement in overall fit. For overall cup accuracy (Figure 3B), all three models produced values in the observed range, but the attention-only and combined models again provided the closer descriptive match. By contrast, the referential-only model tended to misallocate accuracy across the condition subsets. The most informative comparison concerned conditional cup accuracy (Figure 3C). Here the referential-only model clearly overpredicted performance for the 2-side subset, yielding a level of conditional localisation that was not observed empirically. The attention-only model, by construction, remained at the relevant within-side chance baselines for the conditional analysis and therefore captured only the side-first component of performance. The combined model provided the best descriptive compromise, accommodating the strong side-level structure while allowing a limited residual cup-level component. Formal model comparison supported this pattern. The referential-only model showed the weakest fit (log *L* = −660.62, *AIC* = 1327.2, *BIC* = 1340.4), whereas the attention-only model performed better (log *L* = −632.41, *AIC* = 1270.8, *BIC* = 1284.0). The combined model achieved the best log-likelihood and the lowest AIC (log *L* = −627.84, *AIC* = 1267.7), but because it required three additional parameters, its BIC (1294.0) was higher than that of the simpler attention-only model. Thus, the modelling results indicate that most of the structured variance in the present data is already captured by an attentional account, while any additional referential component appears comparatively modest.

### Computational decomposition and generative model comparison

#### Decomposition of exact-cup success

The observed cup-success rate can be decomposed as

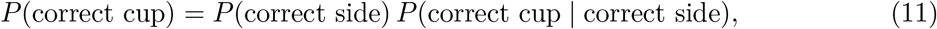

because a correct cup choice requires that the dog first move to the correct side and then select the correct cup within that side. This decomposition clarifies why overall cup accuracy can remain low even when side accuracy is comparatively high. It also isolates the component most relevant to the referential versus attentional contrast, namely *P* (correct cup | correct side).

### Chance predictions under an attentional account

Under a coarse attentional account, the point shifts behaviour toward the indicated side, while within-side selection is random in the absence of exact referent identification. Therefore:

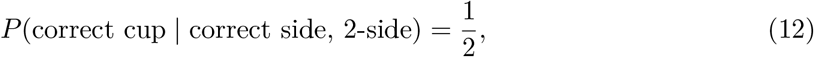

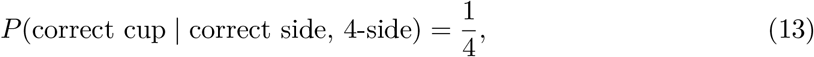

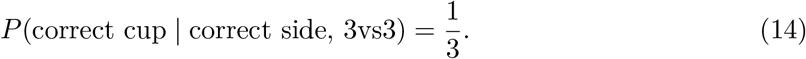

These values are not the baselines for overall cup accuracy. Rather, they are the baselines for *within-side localisation conditional on successful side entry*. Overall cup chance remains 1/6 in a six-cup task.

### Observed pattern relative to the conditional baselines

Comparing the observed values to Eqs. (12)–(14) gives the following pattern:

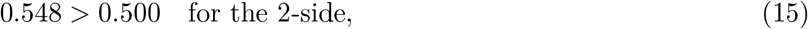

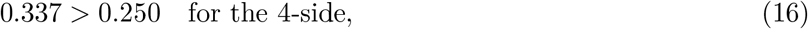

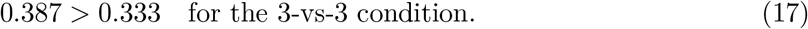

Descriptively, conditional cup accuracy exceeded the relevant within-side chance baselines in all three pointing subsets, although the mixed-effects models did not reveal robust between-group differences once repeated observations and chance offsets were taken into account.

### Interpreting overall and conditional cup accuracy in the unequal arrangement

The unequal 2-vs-4 arrangement is most informative when performance is decomposed into two stages: first, whether the dog reaches the correct side of the arena, and second, whether it identifies the correct cup once that side has been reached. As shown in Eq. (11), overall cup accuracy can be written as the product of side accuracy and conditional cup accuracy. This decomposition is important because the same overall cup accuracy can arise from different combinations of side guidance and within-side localisation. A first reference point for overall cup accuracy is the task-level chance rate of the six-cup task, *P*(correct cup by chance) 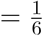, which is the appropriate baseline if choices are treated as globally random over the full array. However, a second and more informative reference point is given by a side-first model in which the gesture guides the dog only to the correct side, after which cup choice within that side is random. Under that two-step account, expected overall cup accuracy is

*P*(correct cup)_side-only_ = *P*(correct side) *× P* (correct cup | correct side, random within side). In the present unequal arrangement, this yields

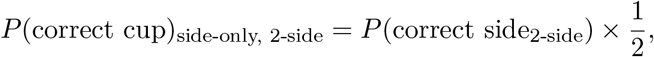

And

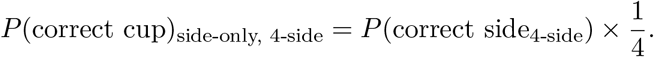

This distinction clarifies the role of Figure 3B. Relative to the global six-cup chance rate of 1/6, overall cup accuracy in both the 2-side and 4-side subsets is above chance. However, Figure 3B alone does not show whether this success reflects exact cup localisation or successful side entry followed by random within-side choice. This question is addressed in Figure 3C, where cup accuracy is evaluated conditional on correct-side entry and compared against the relevant within-side chance baselines given in Eqs. (12) and (13). Under this decomposition, the present pattern is asymmetric, but not in the sense of a simple success on one side and chance performance on the other. On the 2-side, conditional cup accuracy lies only slightly above the relevant within-side chance level of 1/2, whereas on the 4-side it exceeds the lower within-side chance level of 1/4 by a larger descriptive margin. This means that overall cup success cannot be reduced straightforwardly to side guidance followed by random within-side choice, but it also means that the descriptive residual above chance is not strongest where the number of remaining alternatives is smallest.This pattern suggests that the pointing cue affected behaviour at more than one spatial scale. The dominant effect was expressed at the level of side guidance, whereas any additional localisation within the reached side was weaker. Thus, the decomposition supports a layered interpretation, but it does not provide strong evidence for robust object-specific reference. The most conservative conclusion is that dog point-following in the present task was dominated by coarse spatial guidance, with limited residual localisation after correct-side entry.

### Observed grouped outcomes

The grouped counts used for the model comparison are shown in Table 5. Across the three pointing subsets, the strongest descriptive difference again appeared at the level of side guidance. The observed side probabilities were 0.593 for `3vs3`, 0.618 for `2vs4_2side`, and 0.786 for `2vs4_4side`. Overall cup probabilities were lower, at 0.230, 0.338, and 0.265, respectively. Conditional cup accuracy was 0.387 for `3vs3`, 0.548 for `2vs4_2side`, and 0.337 for `2vs4_4side`. Thus, the grouped counts recapitulated the main descriptive pattern already seen in the trial-wise summaries: especially strong side guidance for the 4-side subset, together with more modest exact-cup performance.

**Table 5:** Supplementary trial-level covariate effects in the extended mixed-effects models. Values show omnibus tests for cue side, cue position number, and block after inclusion alongside the main condition factor. Dog identity was included as a random intercept in all models.

| Outcome | Covariate | Test statistic | df | *p* |
| --- | --- | --- | --- | --- |
| Side accuracy | cue_lr | $F = 0.32$ | 1, 586 | .571 |
| Side accuracy | cue_num | $F < 0.001$ | 1, 586 | .997 |
| Side accuracy | block | $F = 1.43$ | 6, 586 | .200 |
| Overall cup accuracy | cue_lr | $F = 0.31$ | 1, 586 | .581 |
| Overall cup accuracy | cue_num | $F = 2.34$ | 1, 586 | .126 |
| Overall cup accuracy | block | $F = 1.05$ | 6, 586 | .394 |
| Within-side localisation | cue_lr | $F = 0.06$ | 1, 408 | .813 |
| Within-side localisation | cue_num | $F = 3.08$ | 1, 408 | .080 |
| Within-side localisation | block | $F = 0.72$ | 6, 408 | .633 |

### Generative model comparison

The model comparison is summarised in Table 6. The referential-only model fit the data worst (log *L* = −660.62, AIC = 1327.2, BIC = 1340.4), indicating that a pure exact-target account did not provide a good description of the grouped outcome structure. The attention-only model performed substantially better (log *L* = −632.41, AIC = 1270.8, BIC = 1284.0), consistent with the strong side-level effects observed in the descriptive and mixed-effects analyses. The combined model achieved the best AIC (log *L* = −627.84, AIC = 1267.7), but because it contained twice as many free parameters its BIC (1294.0) was higher than that of the simpler attention-only model. This pattern suggests that the main structure in the grouped counts is captured by an attentional process, while any additional gain from including a referential component is modest. Put differently, the data do not support a pure referential account, and the principal competition is between a parsimonious attention-only description and a slightly more flexible mixed account.

**Table 6:** Observed grouped outcome counts and derived probabilities for the three pointing subsets used in the generative model comparison.

| Group | Side size | $N$ | Correct cup | Wrong cup, correct side | Wrong side | $P(\text{side})$ | $P(\text{cup})$ |
| --- | --- | --- | --- | --- | --- | --- | --- |
| 3vs3 | 3 | 209 | 48 | 76 | 85 | 0.593 | 0.230 |
| 2vs4_2side | 2 | 204 | 69 | 57 | 78 | 0.618 | 0.338 |
| 2vs4_4side | 4 | 215 | 57 | 112 | 46 | 0.786 | 0.265 |

**Table 7:**
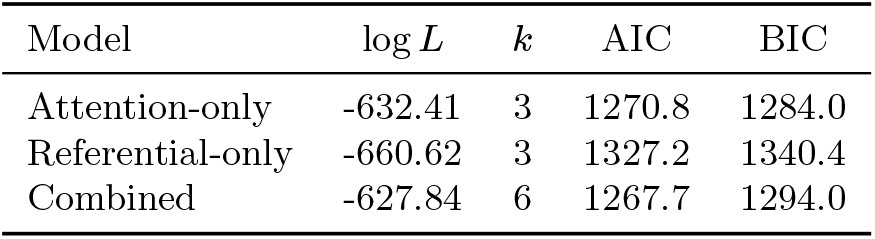
Model comparison for the grouped-outcome generative models. Lower AIC and BIC indicate better fit after penalizing model complexity.

**Table 8:** Parameter estimates for the grouped-outcome generative models.

| Group | $p_{\text{attention}}$ | $q_{\text{referential}}$ | $q_{\text{combined}}$ | $p_{\text{combined}}$ |
| --- | --- | --- | --- | --- |
| 3vs3 | 0.623 | 0.089 | 0.050 | 0.603 |
| 2vs4_2side | 0.646 | 0.225 | 0.062 | 0.623 |
| 2vs4_4side | 0.833 | 0.137 | 0.097 | 0.815 |

### Estimated model parameters and interpretation

The fitted parameters help clarify what the model comparison implies. In the attention-only model, the fitted attention parameters were *p*_attention_= 0.623 for `3vs3`, 0.646 for `2vs4_2side`, and 0.833 for `2vs4_4side`, directly mirroring the observed side-accuracy pattern. Under this model, conditional cup performance is fixed at within-side chance (1/3, 1/2, and 1/4, respectively), so the model necessarily underpredicts the observed conditional cup values. In the referential-only model, the fitted referential parameters were small overall (*q*_referential_ = 0.089, 0.225, and 0.137 for `3vs3`, `2vs4_2side`, and `2vs4_4side`, respectively), and the model fit remained poor. In the combined model, the fitted referential components were smaller still in two of the three subsets (*q* = 0.050, 0.062, and 0.097), whereas the attentional parameters remained comparatively large (*p* = 0.603, 0.623, and 0.815). Thus, when both processes were permitted, the best-fitting solution still assigned most of the explanatory weight to the attentional component, with only a weak residual referential contribution. The fitted models therefore align closely with the main behavioural interpretation developed above. A pure referential account is not supported. A pure attention account captures most of the structure in the data and is preferred by BIC. At the same time, the small AIC advantage of the combined model and the descriptively above-chance conditional cup values are consistent with a layered account in which a dominant attentional process is supplemented by a weaker target-specific component.

## Discussion

The present analysis was designed to test whether an unequal side structure can reveal more about dog point-following than a symmetric multi-cup arrangement alone. A symmetric 3-vs-3 design can already show that side-following and exact cup-following diverge, but it cannot test whether exact localisation scales with the number of alternatives on the indicated side. The unequal 2-vs-4 arrangement adds precisely this leverage. Three main conclusions follow from the present results. First, the clearest condition effect was expressed at the level of *side guidance*. Descriptively, side accuracy was highest when the cued cup was located on the 4-cup side of the unequal arrangement. This pattern was confirmed by the mixed-effects analysis: the effect of group was significant for `side_correct`, and the positive effect was driven specifically by the `2vs4_4side` condition relative to the `3vs3` reference. This effect remained detectable even after the inclusion of `cue_lr`, `cue_num`, and `block` as additional predictors. Thus, the unequal arrangement altered coarse spatial guidance in a systematic way. This interpretation is consistent with earlier work suggesting that dogs often use human pointing at the level of broad directional guidance rather than exact target specification (Dorey et al., 2010; Elgier et al., 2012; Udell et al., 2010; Wynne, 2016; Bowers et al., 2025). In particular, multi-option studies have shown that dogs can follow the general direction of a gesture even when precise localisation remains weak, and this has been taken as evidence that point-following may operate primarily through side- or region-level cue use rather than full object-specific reference (Lakatos et al., 2012; Kaminski and Nitzschner, 2013; Bowers et al., 2025). Second, the pattern for *overall cup accuracy* was weaker and less robust than the corresponding pattern for side guidance. Although exact cup choice was above the relevant unconditional chance levels descriptively, the mixed-effects analyses showed only limited evidence of condition dependence at this level. In the simpler model, the omnibus effect of group reached significance, and this effect was driven by the `2vs4_2side` subset relative to `3vs3`. However, this effect was no longer reliable after the inclusion of cue side, cue position number, and block. Thus, once side selection and within-side localisation were collapsed into a single binary outcome, the unequal arrangement produced at most a modest and comparatively unstable change in exact-cup success across conditions. This suggests that the manipulation primarily affected the level of broad directional guidance rather than robust precise cup identification. Third, and most importantly, the critical within-side analysis did not reveal a comparably strong condition effect. When the analysis was restricted to trials on which the dog had already reached the correct side, and when the condition-specific chance baselines (1/2, 1/4, and 1/3) were taken into account through a logit offset, the effect of group was not significant in the core mixed model and remained non-significant in the more complex extensions. Thus, the unequal arrangement did not yield strong evidence that dogs localised the target cup more precisely in one condition than another once correct-side entry had already been achieved. Taken together, these results support a dissociation between *getting into the correct region* and *selecting the correct cup within that region*, with the former showing the clearest and most robust condition dependence. The strongest effect was observed at the level of side guidance, especially when the correct side was the larger side, whereas cup-level effects were weaker and less stable across model specifications. Conditional cup accuracy was descriptively above the relevant within-side chance baselines in all three pointing subsets, indicating that behaviour was not exhausted by coarse side guidance alone. However, the mixed-effects analyses did not yield robust group differences at this stage.

This conditional analysis clarifies how above-chance cup performance should be interpreted. Overall cup accuracy exceeding the global six-choice chance level does not by itself distinguish between exact cup localisation and successful side entry followed by random within-side choice. Evaluating performance conditional on correct-side entry isolates this second stage of the process. The resulting pattern indicates that pointing primarily improved the probability of reaching the correct side, while any additional localisation within that side remained comparatively weak. A mixed interpretation of this kind is compatible with broader critical discussions of the object-choice literature, in which above-chance responding has been argued to arise from multiple contributing processes rather than from a single, fully referential mechanism. Reviews and meta-analytic work have emphasized that dogs’ success in pointing paradigms is strongly shaped by methodological features and does not, by itself, settle the question of whether dogs represent the gesture as object-specific reference (Udell et al., 2010; Elgier et al., 2012; Clark et al., 2019). The significance of the unequal arrangement lies precisely here. In a symmetric 3-vs-3 design, it can be shown that correct-side performance exceeds exact-cup performance, but the design cannot determine whether exact-cup performance behaves like a side-first search process whose difficulty depends on the number of alternatives within the indicated side. The 2-vs-4 manipulation was intended to test this more specific prediction. The present data indicate that the manipulation clearly influenced side behaviour, but did not produce equally clear changes in within-side cup localisation. This suggests that the principal effect of the gesture is not to specify a particular cup robustly, but to bias movement toward a broad spatial region. The particularly high side accuracy observed when the correct side was the 4-cup side is theoretically informative. A simple “fewer options is easier” account would not predict this pattern at the level of side choice. Instead, the larger side appears to have functioned as a stronger spatial attractor. One plausible interpretation is that the gesture was used first at a coarse spatial scale, such that a larger indicated region was easier to follow as a region. Once that region had been reached, cup choice was not entirely random, suggesting that some target-specific information remained available. On this view, the unequal arrangement reveals two partially opposing tendencies: a large-side attentional advantage that facilitates correct region entry, and a smaller-side localisation advantage that would in principle facilitate exact cup choice once the correct side has been reached. The present data suggest that the first of these tendencies was the more robust one, whereas the second was expressed only as a weaker referentially relevant component. This interpretation also fits with eye-tracking and related process-oriented accounts suggesting that dogs are highly sensitive to the attentional and spatial properties of human communicative displays. If the gesture first shifts attention toward a broad region, then a larger cued side may naturally exert a stronger pull, even when precise object-level localisation remains comparatively limited (Rossi et al., 2014).

At the same time, the present findings do not imply that the gesture acted only as a coarse side cue. Descriptively, conditional cup accuracy was above the relevant within-side chance baselines in some subsets, and the mixed-effects models did not suggest complete absence of signal. Rather, the present pattern suggests that whatever information is extracted from the point is expressed more strongly at the level of coarse directional guidance than at the level of precise referent identification. In this sense, the results align with the broader interpretation that dogs are influenced by human pointing, but not in a way that strongly supports a human-like, object-specific referential reading of the gesture. In this respect, the present results also resonate with studies arguing that dogs may extract more than mere direction from human gestures without thereby demonstrating a fully human-like referential interpretation. For example, dogs’ performance has been shown to vary with the communicative context, the presence of ostensive signals, and the wider informational structure of the task, suggesting that target-specific behaviour may emerge in graded rather than all-or-none form (Scheider et al., 2013; Tauzin et al., 2015).

Viewed through broader theoretical frameworks, the present results suggest a layered form of cognition rather than a strict opposition between “simple” and “complex” mechanisms. The dominant component of behaviour appears to be a coarse attentional process that guides dogs toward the relevant region. However, because cup choice after correct-side entry was not uniformly reducible to within-side chance, the data also suggest a second, more selective component that is sensitive to the specific target. This does not require positing fully human-like referential inference, but it does indicate that the behaviour is not exhausted by attentional capture alone. Theoretical progress may therefore lie in recognising that point-following can recruit multiple levels of processing within the same behavioural sequence. The complementary generative model comparison reinforces this reading. At the grouped-count level, the referential-only model performed worst, whereas the attention-only model captured the main structure of the data substantially better. The combined model achieved the lowest AIC, but only with a small improvement over the attention-only model and with relatively small fitted referential weights. BIC, which penalizes additional parameters more strongly, preferred the simpler attention-only account. Taken together, these results suggest that the data are not well described by a pure exact-target mechanism, but are most consistent with a dominant attentional process and, at most, a modest residual referential contribution.

A further point is that the nuisance covariates added little to the main theoretical picture. Cue side (`cue_lr`) did not exert reliable omnibus effects in the extended models, and cue position number (`cue_num`) was likewise non-significant, although it showed a weak negative tendency in the within-side analyses. Block did not show a significant omnibus effect in any of the three extended models, even though individual block coefficients reached significance in the side-accuracy model. Accordingly, the principal conclusion does not appear to depend on left-right asymmetry, simple block progression, or a strong monotonic position effect.

Exploratory supplementary analyses including selected dog-level metadata variables likewise did not alter the principal interpretation in the present dataset, but they add one theoretically relevant qualification. Training time was not associated with side accuracy, but it was positively associated with both overall cup accuracy and within-side localisation in the reduced combined metadata models. This pattern suggests that training history may contribute specifically to the more precise cup-level components of performance, rather than to the initial side-guidance component. Because these analyses were exploratory and based on a modest sample, this effect should be treated as a secondary individual-difference result rather than as a central explanatory factor. One possible interpretation is that training history modulates the spatial resolution at which dogs use the pointing cue. Dogs with longer training histories may attend more closely to fine-grained spatial features of the experimenter’s cue, such as pointing direction, hand position, gaze shift, or body orientation. This could allow them to use the gesture not only as a coarse side cue, but also as a source of limited information for cup-level localisation. Less trained dogs may still extract the broader directional component of the cue, while relying less on the finer spatial information needed for accurate within-side choice.

Several limitations should nevertheless be acknowledged. First, the present account rests on binary response variables derived from trial outcomes, and the within-side analysis reduces the data to the subset of correct-side trials. Second, although the random-intercept models were stable and the random-slope extension yielded a similar qualitative result, the sample remains modest, and more extensive data would be desirable for estimating subject-specific heterogeneity with greater precision. Third, the current analyses concern behavioural choice structure and do not by themselves determine the cognitive mechanism underlying the observed biases. The present inference is therefore comparative rather than absolute: the data are more readily explained by a coarse attentional account than by a strong referential account. Despite these limitations, the unequal side manipulation appears to provide useful diagnostic value. Its main contribution is not simply to show that correct-cup accuracy is only modestly above chance, but to clarify *where* the pointing effect is expressed. The present results suggest that the effect is expressed most clearly in broad side guidance, especially when the indicated side is larger, whereas exact identification of the specific target cup remains comparatively weak. In this sense, the unequal arrangement extends the logic of the symmetric multi-cup task and provides a more discriminating test of attentional versus referential interpretations of dog point-following. The model comparison reinforced this interpretation. Although the combined model achieved the best descriptive fit in AIC terms, the gain over the simpler attention-only model was modest, and the BIC favoured the attention-only account. At the same time, the referential-only model provided the poorest fit and overpredicted conditional localisation in the 2-side subset. Thus, the modelling results align with the behavioural decomposition in suggesting that the dominant component of performance lies in coarse side guidance, while any additional target-specific component is comparatively weak.

At a broader theoretical level, the present results do not fully settle the long-standing debate between referential and attentional accounts of dog point-following, but they do constrain it more sharply. The data are difficult to reconcile with a purely low-level attentional account in which the gesture does no more than bias approach toward one side, because cup choice after correct-side entry was not wholly reducible to within-side chance. At the same time, the results are also difficult to reconcile with a strong referential account in which the gesture functions primarily as a direct signal of the specific target cup, because the clearest and most robust effects were expressed at the level of broad side guidance rather than exact cup localisation. The most plausible resolution suggested by the present analysis is therefore an intermediate one: pointing appears to function first as a coarse guide to the relevant spatial region, while more precise target-related information contributes only secondarily once that region has been reached. The theoretical progress made here lies not in deciding unambiguously for one classical position over the other, but in showing more clearly how both components may coexist within the same behavioural sequence.

## Author Contributions (CRediT role names)

Jennifer D. Mugleston: Conceptualization, Investigation (experimental testing and data collection), Data curation, Methodology.

Shin-Miau Huang: Conceptualization, Investigation (experimental testing and data collection), Data curation, Methodology.

Christoph D. Dahl: Conceptualization, Investigation (experimental testing and data collection), Data curation, Methodology, Formal analysis, Software, Visualization, Supervision, Project administration, Writing – original draft, Writing – review and editing, Funding acquisition, Resources.

## Funding

This research was funded by the National Science and Technology Council (NSTC), Taiwan, under grant number 114-2410-H-038-045, and by the Higher Education Sprout Project of the Ministry of Education (MOE), Taiwan, both awarded to C.D.D.

## Institutional Review

All procedures were reviewed and approved by the Taipei Medical University Institutional Animal Care and Use Committee (IACUC; approval number: SHLAC2023-0072). The study involved non-invasive behavioural testing of privately owned dogs. Owners provided informed consent, and all participation was voluntary.

## Data Availability

The anonymised trial-level data, derived analysis files, MATLAB scripts used for preprocessing, statistical analyses, model comparison and figure generation, and a README file describing the folder structure and workflow are made available in a public GitHub repository: https://github.com/ChristophDahl/dog-pointing.

## Conflict of interest

The authors declare no conflict of interest. The funders had no role in the design of the study; in the collection, analysis, or interpretation of data; in the writing of the manuscript; or in the decision to publish the results.

## Notes

https://github.com/ChristophDahl/dog-pointing

### Competing Interest Statement

The authors have declared no competing interest.

### Summary of Updates

Adding caption for table 3, which was missing in the previous version.

